# Identification of critical genes and biological signaling for metformin treated liver cancer

**DOI:** 10.1101/2021.12.29.474467

**Authors:** Tingting Zhang, Hongmei Guo, Letian Wang, Mengyao Wang, Hanming Gu

**Author notes:** Corresponding author: Hanming Gu, SHU-UTS SILC School, Shanghai University, Shanghai, China.

## Abstract

Liver cancer is a leading source of cancer-related mortality in the world. A number of studies have shown the correlation of metformin treatment with a decrease in cancer risk. However, the relevant molecules and mechanisms are not clear during the treatment. In this study, our aim is to identify the significant molecules and signaling pathways in the treatment of metformin in liver cancer cells by analyzing the RNA sequence. The GSE190076 dataset was created by performing the Illumina NovaSeq 6000 (Homo sapiens). The KEGG and GO analyses indicated that DNA synthesis and cell cycle are the main processes during the treatment of metformin. Moreover, we determined numerous genes including RRM2, CDC6, CDC45, UHRF1, ASF1B, ZWINT, PCNA, ASPM, MYC, and TK1 by using the PPI network. Therefore, our study may guide the clinical work on the treatment of liver cancer by using metformin.

## Introduction

Primary liver cancer is an aggressive tumor that is one of the most common cancers in the world^1^. In the United States, the annual incidence of liver cancer is increasing despite the advanced treatment, and liver cancer remains one of the most difficult cancers to cure^2^. There are various treatments for liver cancer, which are selected according to the complex interplay of tumor stage and patient’s status^3^.

Metformin is an essential drug for the treatment of diabetes, which is the most commonly used oral drug and is recommended as the first-line therapy for T2D patients^4^. Besides diabetes, metformin is also used in polycystic ovary diseases, diabetic nephropathy, and cardiovascular complications^5^. Most importantly, metformin indicated the anti-cancer functions through both indirect and direct effects^6^. The indirect mechanisms contribute to the regulation of blood glucose and insulin, which could affect cancer cell survival^7^. Metformin directly inhibits the NF-κB activity to further activate the immune response to cancer cells^8^. For example, metformin enhanced the effectiveness of the anti-cancer vaccine that was regulated by the activation of T cells^9^.

Here, our present studies determined the DEGs and signaling pathways of metformin in liver cancer by analyzing the RNA-seq data. We figured out several functional DEGs and processes and created the protein-protein interaction (PPI) network. It is possible that our understanding of the anti-cancer properties of metformin will result in advances in the treatment of liver cancer.

## Methods

### Data resources

The data (GSE190076) was created by using Illumina NovaSeq 6000 (Homo sapiens) (Hebei University, Number 180, Wusi east street, Baoding, Hebei Province, China). The analyzed dataset includes three groups of control liver cancer cells and three groups of liver cancer cells treated by metformin.

### Data acquisition and preprocessing

The data were processed by the R package as described^10-14^. A classical t-test was performed to identify DEGs with P< 0.01 and fold change ≥ 1 as being statistically significant.

### The Kyoto Encyclopedia of Genes and Genomes (KEGG) and Gene Ontology (GO) analyses

The KEGG and GO analyses were performed from the R package (clusterProfiler) and Reactome (https://reactome.org/). We set the P<.05 and gene counts >10 as the statistically significant cutoff.

### Protein-protein interaction (PPI) analysis

The PPI was constructed by using the Molecular Complex Detection (MCODE). The biological processes analysis was performed by Reactome, and P<0.05 was considered as the statistically significant cutoff.

## Results

### Identification of DEGs of metformin treated liver cancer cells

To understand the mechanism of the effects of metformin on liver cancers, we analyzed the RNA-seq data from the metformin treated liver cancer cells. A total of 125 genes were identified with the threshold of P < 0.001. The top up-and down-regulated genes in metformin treated liver cancer cells were identified by the heatmap and volcano plot (Figure 1). The top ten DEGs were listed in Table 1.

**Table 1.**

| **Entrez gene** | **Gene symbol** | **Fold-change** | **Regulation** |
| --- | --- | --- | --- |
| Top 10 down-regulated DEGs |  |  |  |
| 6236 | RRAD | -5.641311351 | Down |
| 80201 | HKDC1 | -5.331256314 | Down |
| 4688 | NCF2 | -5.019051335 | Down |
| 343 | AQP8 | -4.538374629 | Down |
| 2331 | FMOD | -4.360346053 | Down |
| 119395 | CALHM3 | -4.314311307 | Down |
| 9229 | DLGAP1 | -4.076112 | Down |
| 5649 | RELN | -4.068611599 | Down |
| 153222 | CREBRF | -3.688866916 | Down |
| 4495 | MT1G | -3.639888941 | Down |
| Top 10 up-regulated DEGs |  |  |  |
| 4973 | OLR1 | 4.670169784 | up |
| 374393 | FAM111B | 4.38801419 | up |
| 4091 | SMAD6 | 4.369002101 | up |
| 29128 | UHRF1 | 4.290715153 | up |
| 6241 | RRM2 | 3.73058798 | up |
| 9502 | XAGE2 | 3.67535914 | up |
| 9415 | FADS2 | 3.423215343 | up |
| 65263 | PYCR3 | 3.392965269 | up |
| 54058 | C21orf58 | 3.332096934 | up |
| 147011 | PROCA1 | 3.285038179 | up |

**Figure 1.**
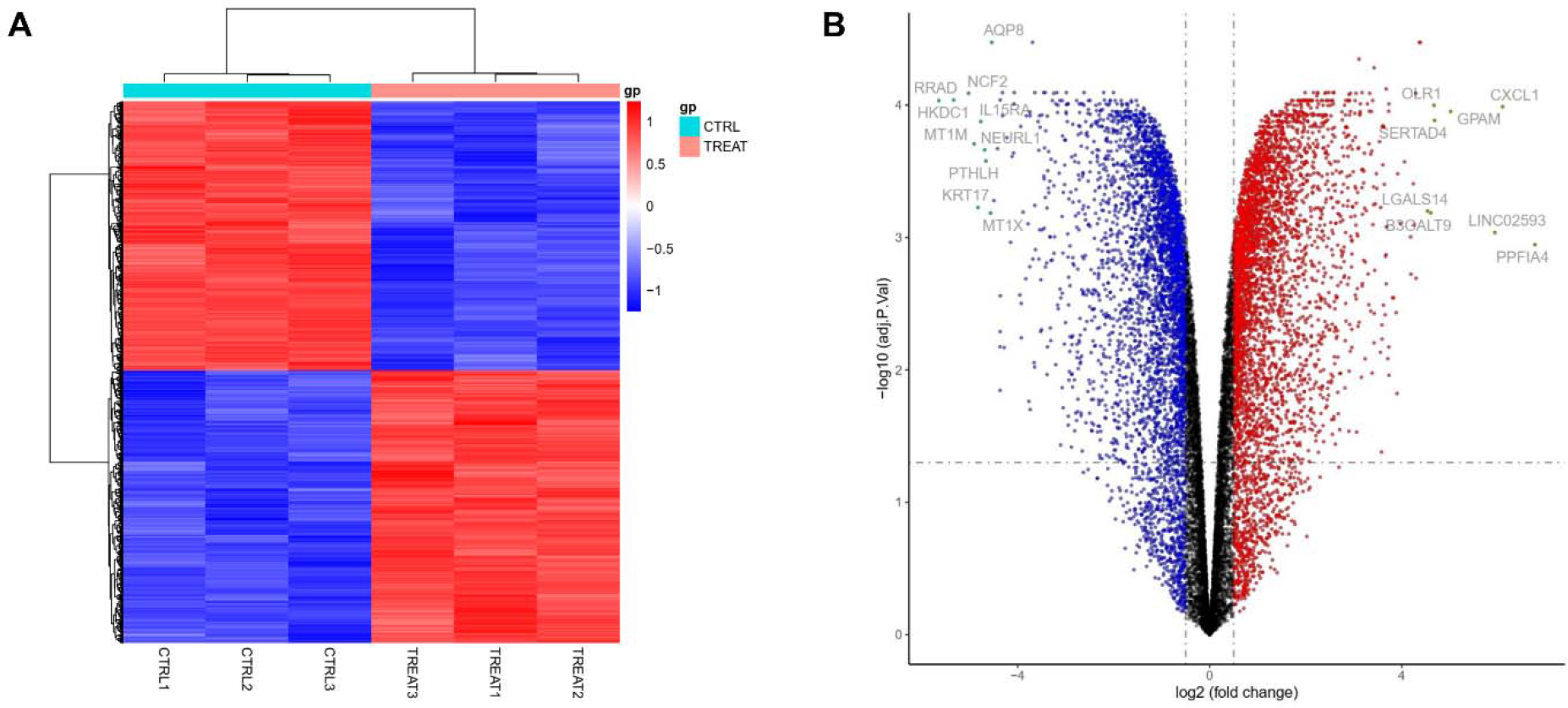
Heatmap and volcano plot were obtained between the control (CTRL) and metformin treated (TREAT) liver cancer cells. (A) Heatmap of significant DEGs. Significant DEGs (P < 0.01) were used to create the heatmap. (B) Volcano plot for DEGs between the control and metformin treated liver cancer cells. The most significantly changed genes are highlighted by grey dots.

### Enrichment analysis of DEGs in the metformin treated liver cancer cells

To further understand the potential biological functions of the DEGs, we introduced the KEGG and GO analyses (Figure 2). The significant items of KEGG were selected, including “Cell cycle”, “Cellular senescence”, “TGF−beta signaling pathway”, “DNA replication”, and “Fanconi anemia pathway”, and “Mismatch repair”. We also identified the top ten biological processes of GO, including “Regulation of mitotic cell cycle”, “Nuclear division”, “Chromosome segregation”, “Negative regulation of cell cycle”, “DNA replication”, “Nuclear chromosome segregation”, “Sister chromatid segregation”, “Mitotic sister chromatid segregation”, “DNA−dependent DNA replication”, and “Cell cycle checkpoint signaling”. We then identified the top ten cellular components of GO, including “chromosomal region”, “spindle”, “condensed chromosome”, “chromosome, centromeric region”, “condensed chromosome, centromeric region”, “mitotic spindle”, “kinetochore”, “spindle midzone”, “DNA replication preinitiation complex”, and “CMG complex”. We identified the top ten molecular functions of GO, including “ubiquitin−like protein transferase activity”, “ubiquitin−like protein ligase activity”, “catalytic activity, acting on DNA”, “histone binding”, “helicase activity”, “single−stranded DNA binding”, “ATP−dependent activity, acting on DNA”, “DNA helicase activity”, “single−stranded DNA helicase activity”, and “DNA replication origin binding”.

**Figure 2.**
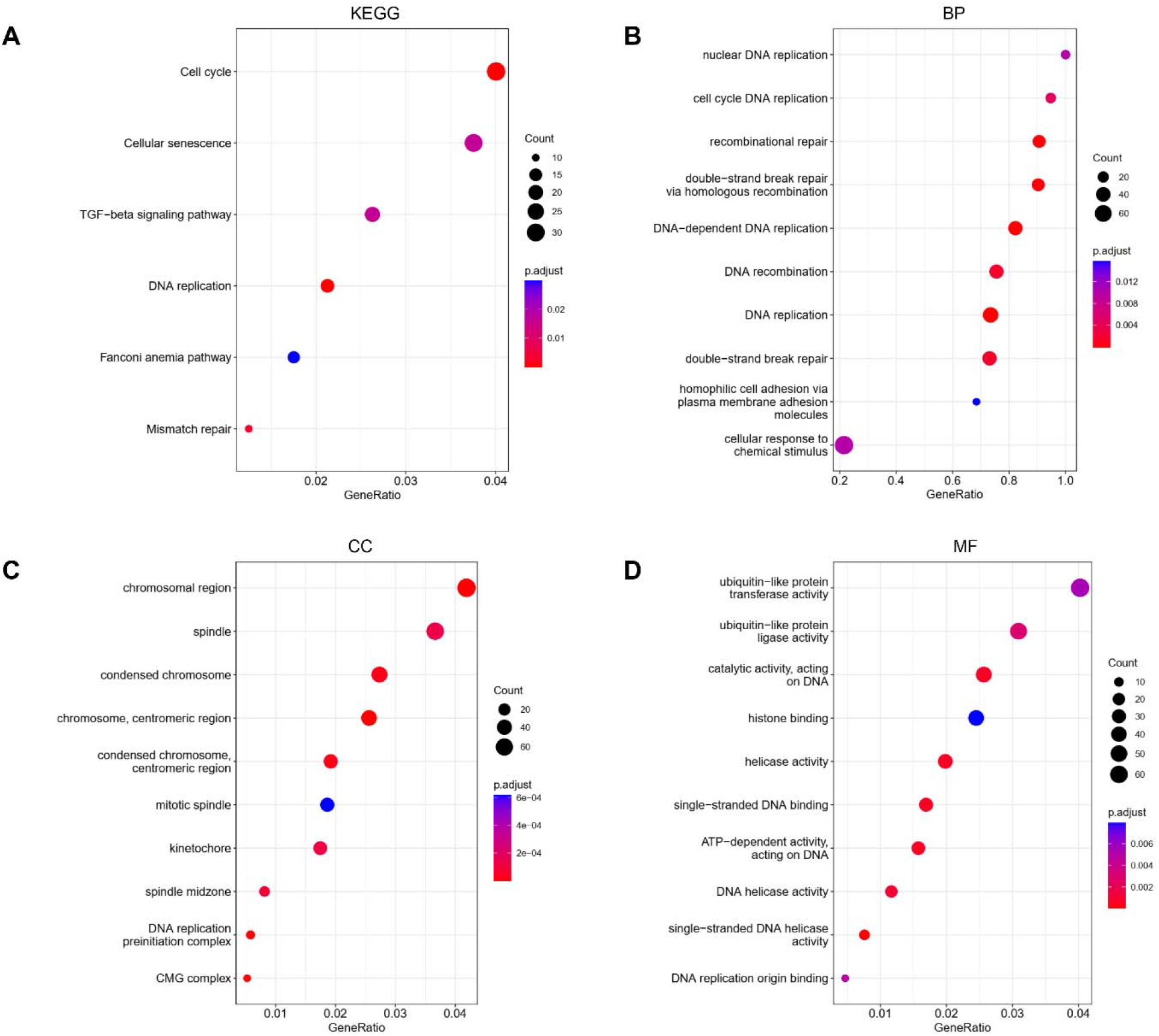
KEGG and GO analyses of DEGs between the control and metformin treated liver cancer cells. (A) KEGG analysis, (B) Biological processes, (C) Cellular components, (D) Molecular functions.

### Construction of PPI network in the metformin treated liver cancer cells

To further identify the potential relationship of the DEGs, we constructed the PPI network by the String and Cytoscope software. The PPI network was created by using 116 nodes and 164 edges (combined score > 0.4). The top ten genes were selected with the highest degree scores (Table 2). The String network and top clusters were shown in Figure 3. We further analyzed the PPI genes and DEGs by Reactome map (Figure 4) and selected the top ten potential biological functions including “G1/S-Specific Transcription”, “Mitotic G1 phase and G1/S transition”, “G1/S Transition”, “Cell Cycle, Mitotic”, “Cell Cycle”, “Transcription of E2F targets under negative control by DREAM complex”, “Transcriptional Regulation by E2F6”, “G0 and Early G1”, and “Activation of ATR in response to replication stress” (Supplemental Table S1).

**Table 2.** Top ten genes demonstrated by connectivity degree in the PPI network.

| **Gene symbol** | **Gene title** | **Degree** |
| --- | --- | --- |
| RRM2 | Ribonucleotide Reductase Regulatory Subunit M2 | 18 |
| CDC6 | Cell Division Cycle 6 | 17 |
| CDC45 | Cell Division Cycle 45 | 17 |
| UHRF1 | Ubiquitin Like With PHD And Ring Finger Domains 1 | 16 |
| ASF1B | Anti-Silencing Function 1B Histone Chaperone | 15 |
| ZWINT | ZW10 Interacting Kinetochore Protein | 15 |
| PCNA | Proliferating Cell Nuclear Antigen | 15 |
| ASPM | Assembly Factor For Spindle Microtubules | 14 |
| MYC | MYC Proto-Oncogene, Bhlh Transcription Factor | 13 |
| TK1 | Thymidine Kinase 1 | 12 |

**Figure 3.**
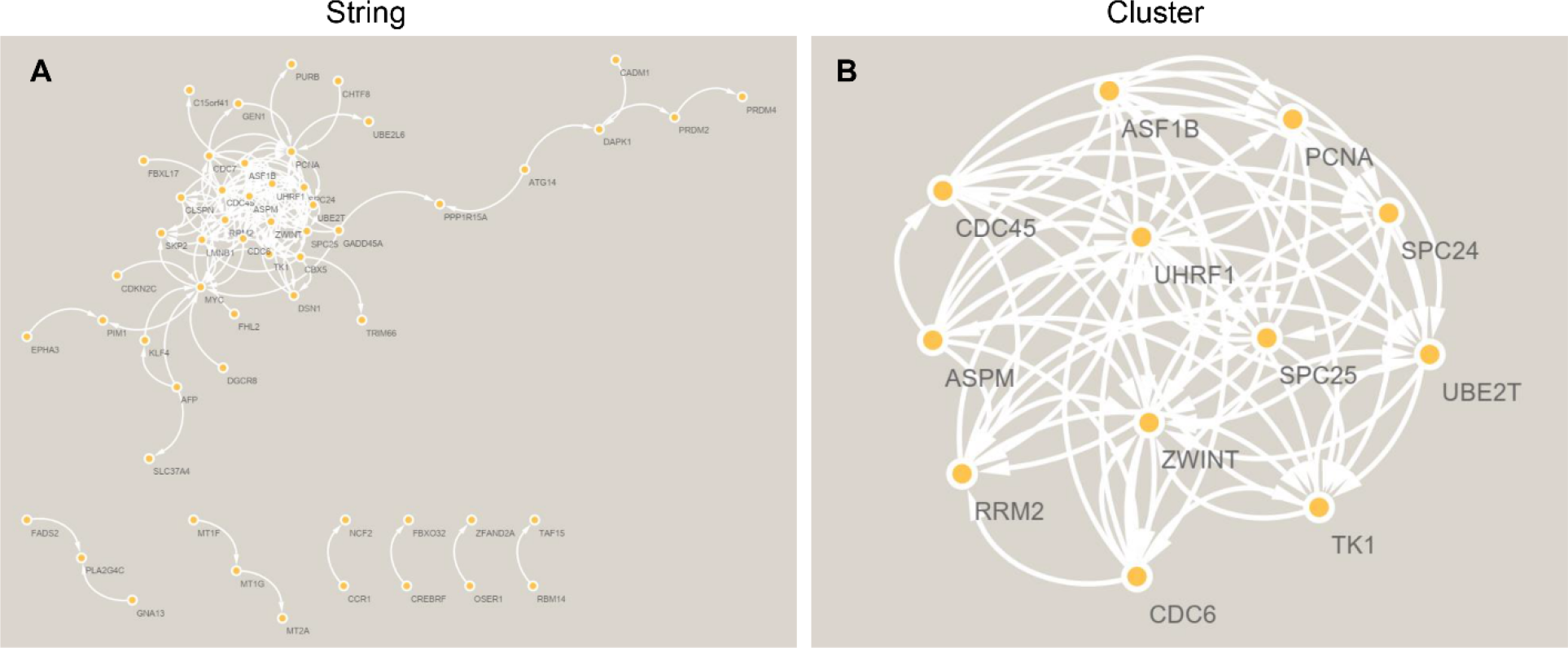
The String and PPI network analyses of DEGs between the control and metformin treated liver cancer cells. String (A) and the first cluster (B) were constructed by MCODE and Cytoscape.

**Figure 4.**
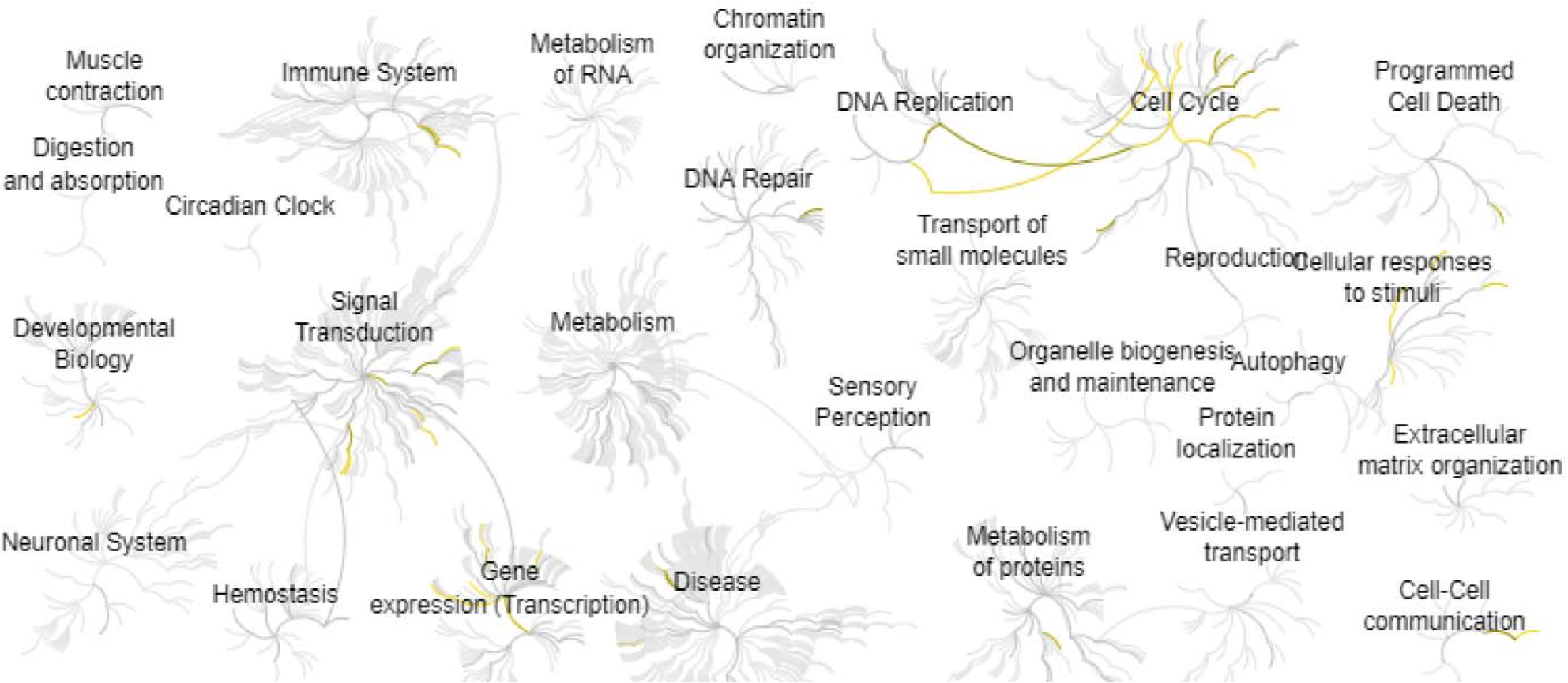
Reactome map representation of the significant biological processes between the control and metformin treated liver cancer cells.

## Discussion

Metformin can inhibit hepatic gluconeogenesis and promote glucose uptake in muscles, and it also can improve the prognosis of cancer patients and block the progression of tumor^6, 15^. Thus, metformin is currently being investigated for new anti-cancer drugs^16^.

Metformin is a key regulator in liver cancer by mediating the DNA replication and cell cycle through the KEGG and GO analyses. Interestingly, Sung-Hee Kim et al found metformin enhances chemo-sensitivity by decreasing DNA replication proteins in colorectal cancer cell^17^. Krisztina Kisfalvi et al found metformin disturbs the relationship between G protein-coupled receptor and insulin receptor signaling systems to further prevent cancer growth^18^. As key signaling regulators, GPCR and RGS proteins are involved in a variety of human diseases including cancer, arthritis, bone diseases, and heart diseases^19-29^. It was found that low doses of metformin inhibited the DNA synthesis and tumor growth by insulin and GPCR agonists^18^. James Sinnett-Smith et al found metformin can inhibit the activation of mTORC1, DNA synthesis in pancreatic cancer by affecting the AMPK signaling^30^. Metformin is capable to inhibit the proliferation of multiple myeloma cells by the inhibition of G0/G1 cell cycle arrest but not apoptosis^31^. Xiaojia Zhou also found metformin restrains the cell proliferation in SKM-1 cells through AMPK-regulated cell cycle arrest^32^. The study by Takuma Yamashita showed metformin prohibits the proliferation in gallbladder adenocarcinoma cell lines including NOZ, TGBC14TKB, and TGBC24TKB, and inhibits the cell cycle^33^.

Besides the important biological processes, we also identified a number of key genes that involve in the treatment of liver cancer with metformin. Yueyue Yang et al found BRM2 is a liver cancer biomarker that indicates specifically increased levels in liver cancer and restraints ferroptosis by triggering GSH synthesis^34^. The high expression of RRM2 was found in 81% of patients with liver cancer and the expression of RRM2 was associated with viral etiology and liver cirrhosis, which may be a biological marker for poor prognosis of liver cancer^35^. CDC6 was identified as a therapeutic target of miR-215 in liver cancer^36^. Moreover, CDC6 is an essential player for DNA replication, which is related to the risk for liver cancer^37^. The circadian clocks and downstream targets are related to a variety of physiological and pathophysiological processes such as cell proliferation, cell differentiation, cell metabolism, immune, and secretion^38-47^. Jorge Fung-Uceda et al found the clock component TOC1 can inhibit DNA replication by binding the CDC6 promoter^48^. Bioinformatic analysis showed the CDC45 gene is closely associated with liver cancer and patient outcomes^49^. Sheng-Ming Wu et al found the knockdown of UHRF1 can inhibit the progression of liver cancer^50^. ASF1A is considered as a prognostic marker, which involves in the progression and development of specific cancers^51^. ZWINT plays key roles in the mitotic checkpoint, which can predict the pathogenesis and prognosis of liver cancer^52^. PCNA is identified as a potential marker for early hepatocellular carcinoma^53^. ASPM is a biological marker for vascular invasion and poor prognosis of liver cancer^54^. Kyu Yun Jang et al found SITR1 and MYC drive liver tumor cell survival and predict liver cancer^55^. Serological TK1 is a critical proliferation biomarker for the early discovery of pre-malignancies^56^.

In conclusion, our study showed the significant impact of metformin in liver cancer. The DNA replication and cell cycle are the major signaling pathways involved in the treatment of liver cancer with metformin. Therefore, this study may be beneficial in the treatment of hepatocellular carcinoma.

## Author Contributions

Tingting Zhang, Hongmei Guo, Letian Wang, Mengyao Wang: Methodology and Writing. Hanming Gu: Conceptualization, Writing- Reviewing and Editing.

